# A KIF1C-CNBP motor-adaptor complex for trafficking mRNAs to cell protrusions

**DOI:** 10.1101/2024.06.26.600878

**Authors:** Konstadinos Moissoglu, Tianhong Wang, Alexander N. Gasparski, Michael Stueland, Elliott L. Paine, Lisa Jenkins, Stavroula Mili

## Abstract

mRNA localization to subcellular compartments is a widely used mechanism that functionally contributes to numerous processes. mRNA targeting can be achieved upon recognition of RNA cargo by molecular motors. However, our molecular understanding of how this is accomplished is limited, especially in higher organisms. We focus on a pathway that targets mRNAs to peripheral protrusions of mammalian cells and is important for cell migration. Trafficking occurs through active transport on microtubules, mediated by the KIF1C kinesin. Here, we identify the RNA-binding protein CNBP, as a factor required for mRNA localization to protrusions. CNBP binds directly to GA-rich sequences in the 3’UTR of protrusion targeted mRNAs. CNBP also interacts with KIF1C and is required for KIF1C recruitment to mRNAs and for their trafficking on microtubules to the periphery. This work provides a molecular mechanism for KIF1C recruitment to mRNA cargo and reveals a motor-adaptor complex for mRNA transport to cell protrusions.

## Introduction

Localization of mRNAs to specific subcellular compartments is a widely used mechanism that critically impacts the spatio-temporal regulation of gene expression and affects various functional outcomes ^1–3^. Biological processes that rely on localization of mRNAs include cell fate determination, embryonic patterning, neuronal outgrowth, synaptic plasticity, and cell migration ^3–6^. Accordingly, mRNA targeting and local regulation is important for ensuring proper organismal development and is deregulated in a variety of neurodegenerative and neuromuscular disorders ^7–9^.

A variety of mechanisms have been described that can lead to asymmetric mRNA distributions. These include random diffusion coupled with local entrapment, transcript degradation coupled to localized protection, and active transport to specific destinations ^9–11^. The latter often occurs along cytoskeletal elements and relies on the action of molecular motors, such as kinesins, dynein, and myosin ^12^. Individual mRNA molecules can serve as cargo of molecular motors, or they can co-assemble into RNA granules for the coordinated transport of multiple mRNAs ^13,14^. Alternatively, hitchhiking of RNAs onto endosomes or lysosomes can support long-range RNA movements ^15–17^.

These various trafficking mechanisms generally involve cis-acting RNA elements that are either necessary or sufficient to confer a specific distribution pattern and are usually found in the 3’ UTRs of localized transcripts. Such localization elements have been sporadically identified for individual transcripts ^18^. Recent studies geared to identify such sequences in a high-throughput manner, have additionally revealed shorter or longer regulatory elements involved in mRNA targeting ^19–21^. How these RNA elements mediate interaction of an mRNA with the machinery that mediates local accumulation is not well understood and detailed molecular understanding is available for only a few cases.

For example, in yeast, localization elements within the *Ash1* mRNA are recognized by the She2p RNA-binding protein, which recruits the type V myosin motor, Myo4p, through the She3p adaptor ^22^. In another case, the Drosophila-specific RNA-binding protein Egalitarian (Egl) links various localized RNA cargoes to the dynein motor. Egl binds to RNA stem loops that mediate polarized transport and additionally associates with the dynein adaptor BICD2 and the dynein light chain subunit of the dynein motor complex ^23^. These components are sufficient to support directed RNA transport in *in vitro* reconstitution experiments, thus defining a minimal transport-competent complex for directed transport to the minus ends of microtubules ^24,25^. A variety of other RNA cargoes rely on kinesin motors for trafficking towards the plus ends of microtubules. In the case of the Drosophila oskar mRNA, the atypical, RNA-binding tropomyosin (Tm1-I/C; aTm1) serves to stabilize the interaction of kinesin heavy chain (KHC) with oskar mRNA ^26,27^ and regulates kinesin activity to allow coordination with dynein-mediated transport at different stages of oocyte maturation ^28^. While a number of other RNA-binding proteins have been implicated in transport events, the exact RNA signals recognized and/or the links to molecular motors have been less well defined, especially in mammalian systems ^29–33^.

In mammalian mesenchymal cells, a robust localization pathway targets mRNAs to peripheral protrusive regions ^34,35^. mRNAs targeted to cell protrusions encode regulators of cell migration and their local translation in peripheral regions ensures efficient cell movement and invasion. Specifically, local protein synthesis promotes co-translational interactions of the nascent proteins that favor promigratory phenotypes ^36,37^. As described in other cases, the regulatory information directing protrusion mRNA targeting is found within the 3’UTRs, which are sufficient to direct peripheral localization of otherwise diffuse mRNAs. Moreover, specific GA-rich regions have critical roles, and interfering with or deleting such GA-rich regions is sufficient to disrupt peripheral localization and perturb cell movement in various systems ^36–39^. Localization to the periphery requires the microtubule cytoskeleton and in particular a subset of stable, detyrosinated microtubules ^35,40,41^. Active trafficking on microtubules is mediated though the kinesin-3 family member KIF1C. KIF1C co-traffics with individual mRNAs along linear paths and is required for their localization, suggesting that it is the main kinesin motor supporting this localization pathway ^42^. Nevertheless, how KIF1C recognizes and is recruited to its RNA cargo has been unclear.

Here, we identify the RNA-binding protein CNBP as the factor recognizing GA-rich regions of protrusion-targeted mRNAs. We demonstrate that CNBP participates in this localization pathway through interacting with KIF1C and serving as an adaptor that recruits the motor to RNA cargo. Our data present a novel motor-adaptor complex that supports trafficking of mRNAs to cellular protrusions.

## Results

### Identification of proteins binding to localization sequences of protrusion-localized mRNAs

Several protrusion-localized mRNAs are targeted to peripheral subcellular locations through associating with the KIF1C kinesin ^42^. Sequences within the 3’UTRs are necessary and sufficient for peripheral targeting through this mechanism ^36,37,39^. We sought to identify factors that connect these mRNAs to the KIF1C motor. For this, we carried out an unbiased mass spectrometry identification of proteins that bind to the 3’UTR of such mRNAs. To narrow down on proteins that might be relevant to the localization pathway, we identified proteins that bind to the full-length mouse *Pkp4* 3’UTR or to 3’UTR fragments that have been shown in prior studies to retain, or not, localization activity ^35^ (Fig. 1A; Pkp4-A and Pkp4-B respectively). These RNA fragments were generated through *in vitro* transcription and were designed to additionally contain at their 3’end a BoxB hairpin sequence from the λ bacteriophage. The BoxB sequence is recognized by a peptide of the λ N protein, which when fused to GST allows immobilization of BoxB-containing RNAs on glutathione beads (Fig. 1A). Incubation with cellular lysates can then allow pulldown of proteins that bind to specific RNAs.

**Figure 1:**
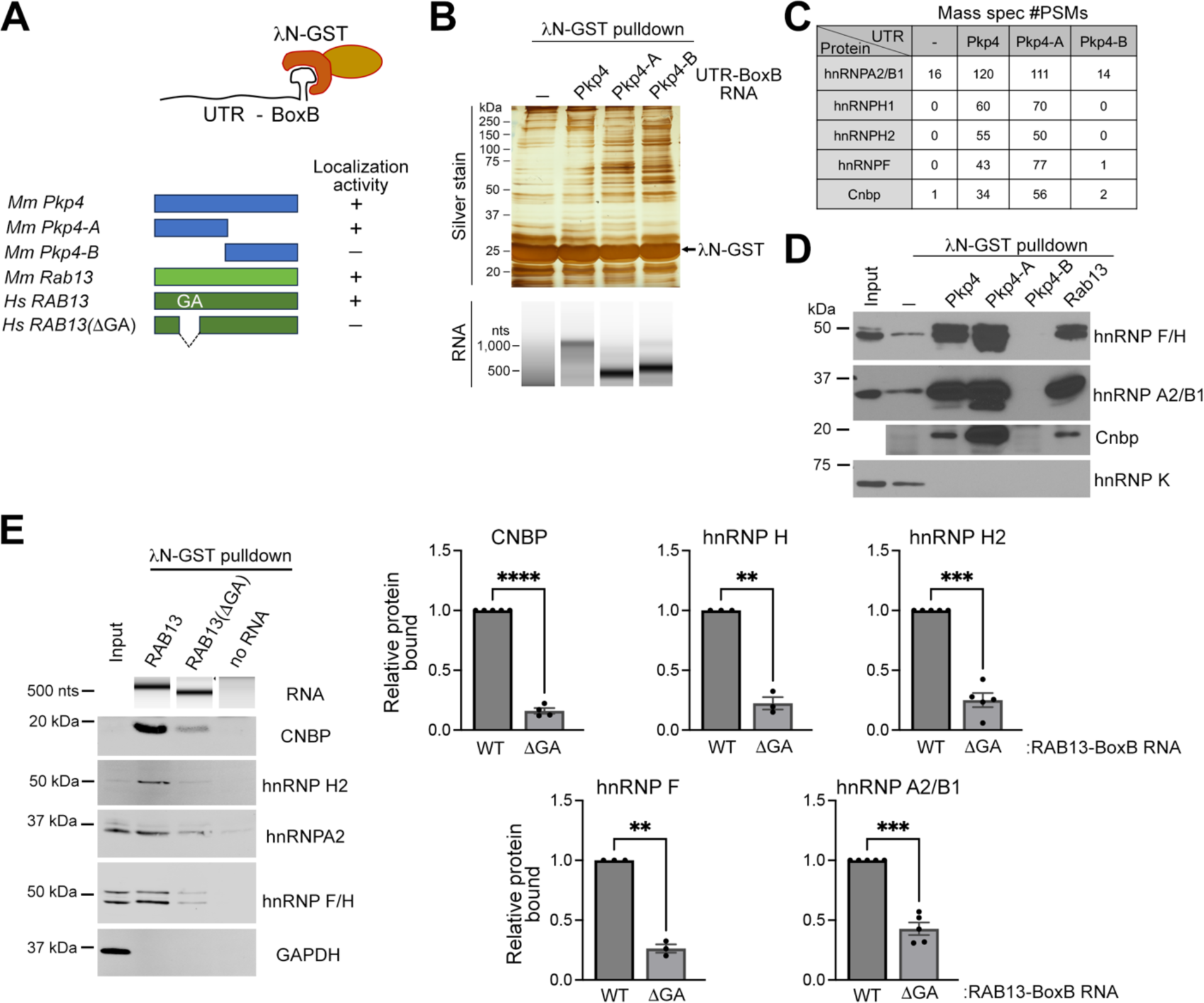
Identification of proteins binding to localization regions of protrusion-localized mRNAs. **(A)** Schematic of BoxB-containing UTR fragments used in λN-GST pulldown assays. Localization activity is indicated based on prior reports (refs ^35,36^). **(B)** Proteins and RNAs recovered after pulldown with the indicated UTRs analyzed through silver staining and TapeStation, respectively. **(C)** Mass spec identification of RBPs preferentially bound to localization-competent Pkp4 UTR fragments. **(D)** Western blot analysis of the indicated proteins in RNA pulldown samples with mouse Pkp4 or Rab13 UTRs. **(E)** RNA pulldown samples with human RAB13 UTRs were analyzed on TapeStation to detect recovered RNAs, or by Western blot to detect the indicated proteins. Graphs indicate quantifications of bound protein to the wild type (WT) or truncated (ΔGA) UTRs. n=3-5. Error bars: SEM. p-values: **<0.01, ***<0.001, ****<0.0001, by paired t-test.

Multiple proteins that specifically bound to *Pkp4* RNA fragments could be visualized by silver staining, despite a high degree of background binding even in the absence of any immobilized RNA (Fig. 1B). These were further identified by mass spectrometry (Fig. 1C and Table S1). We focused on proteins that were preferentially associated with the localization-competent UTR sequences (Pkp4 and Pkp4-A) over either background binding or binding to sequences that do not support localization (Pkp4-B) (Fig. 1C and Table S1). Several RNA-binding proteins (RBPs) were identified that exhibited these characteristics, including hnRNPA2/B1, hnRNPH1, hnRNPH2, hnRNPF and CNBP. To validate the identification of these RBPs, similar pulldowns were performed, and the recovered proteins were analyzed by immunoblotting. Indeed, all the candidate proteins were specifically enriched in pulldowns of localization-competent Pkp4 RNA fragments (Fig. 1D). Other RBPs, such as hnRNPK, were not detected in pulldowns with any Pkp4 RNA sequences, underscoring the specificity of the identified interactions.

Multiple mRNAs are targeted to protrusions through the same KIF1C-dependent mechanism. This localization pathway is also conserved across species ^36,39^. We thus extended our analysis to assess whether the identified RBPs also associated with the 3’UTR of another protrusion-localized mRNA, the mouse *Rab13* mRNA. To further explore conservation among species, we performed pull-down assays using the 3’UTR of the human *RAB13* mRNA and cell lysates from the human MDA-MB-231 cell line. Indeed, we found that the same complement of RBPs associated with the mouse *Rab13* 3’UTR (Fig. 1D), and that the same interactions were conserved in a human system (Fig. 1E; human *RAB13* 3’UTR).

In several cases, localization to protrusions relies on GA-rich regions within the 3’UTRs of targeted mRNAs ^36,37,39,43^. In the case of the human *RAB13* mRNA, when a specific ∼50 nt GA-rich region within the 3’ UTR is deleted, or when its function is interfered with using antisense oligonucleotides, peripheral *RAB13* mRNA localization is impaired ^36^. Therefore, to further explore the potential involvement of the identified RBPs to the localization mechanism, we tested whether the *RAB13* GA-rich sequence is important for their binding. For this, we generated a truncated human *RAB13* 3’UTR fragment missing the GA-rich sequence and performed pulldown assays using human cell lysates. Interestingly, all identified RBPs exhibited significantly reduced binding to the truncated *RAB13* 3’UTR compared to the full-length, wild type (WT) counterpart (Fig. 1E). Taken together, these results indicate that RNA sequences that are necessary and sufficient for mRNA protrusion localization specifically associate with a group of RBPs, making these RBPs candidate participants in the localization mechanism.

### CNBP is required for localization of mRNAs to cytoplasmic protrusions

To evaluate the role of these RBPs in mRNA localization at protrusions, we knocked down their expression in mouse NIH/3T3 cells using siRNAs. Transient knockdown led to a significant decrease of the corresponding RBPs (Fig. S1A). To assess whether this affected mRNA localization to protrusions, we visualized the distribution of protrusion-localized mRNAs (*Rab13*, *Net1*) by fluorescence in situ hybridization (FISH) and quantified it using a previously described Peripheral Distribution Index (PDI) ^44^ (Fig. 2A and S1B). Briefly, values of PDI around 1 denote a random distribution, while values greater or smaller than 1 correspond to a peripheral or perinuclear bias, respectively ^44^ (Fig. 2A). Interestingly, only knockdown of CNBP significantly reduced the peripheral targeting of both *Rab13* and *Net1* mRNAs, indicated by a reduced PDI index, while knockdown of hnRNPA2, hnRNPH1 or hnRNPH2 (individually or in combination) did not significantly or consistently affect peripheral mRNA targeting (Fig. S1B).

**Figure 2:**
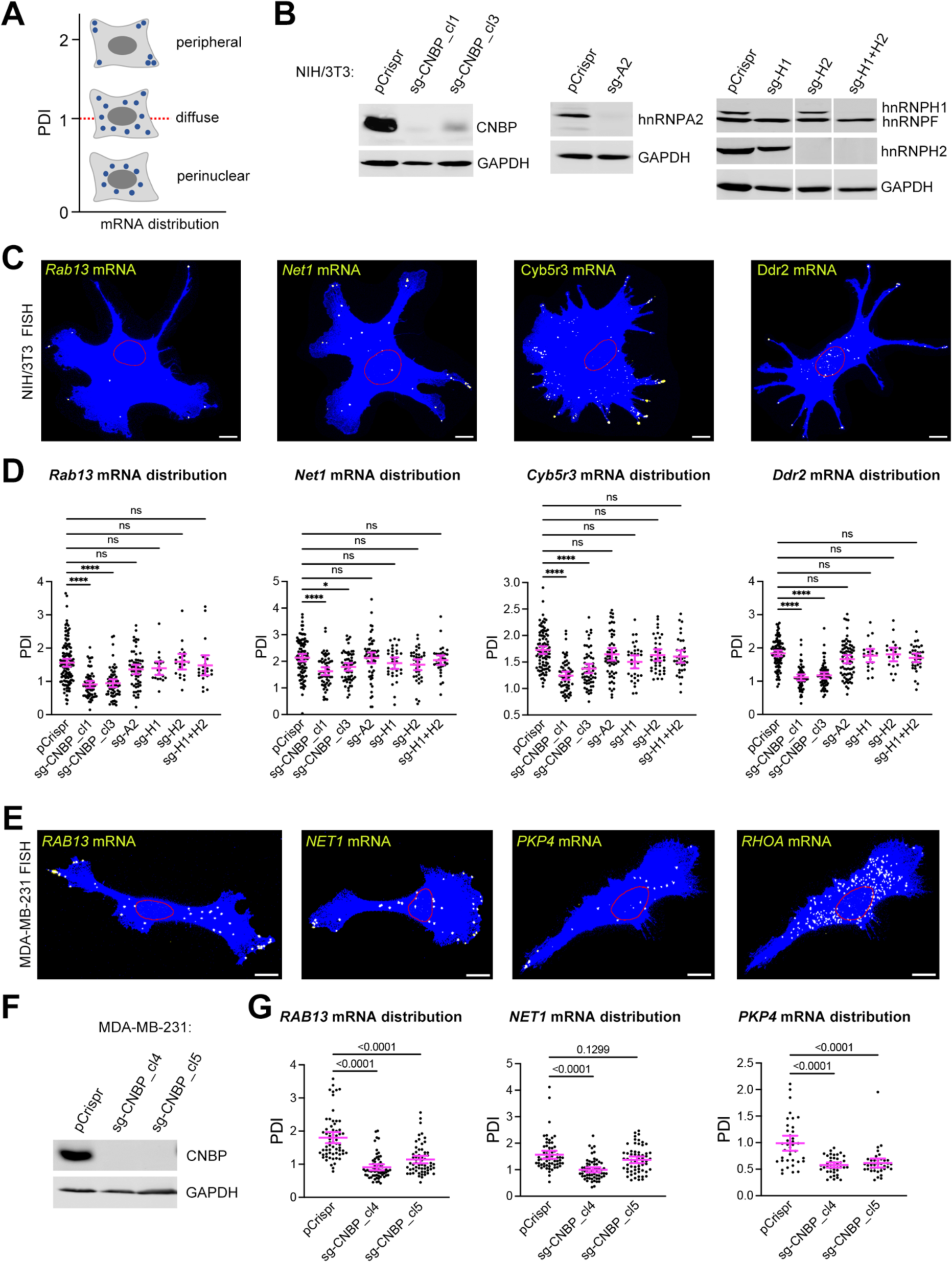
CNBP is required for localization of mRNAs to cytoplasmic protrusions. **(A)** Schematic depicting quantification of RNA distributions through PDI metric. Higher PDI values indicate more peripheral RNA distribution in a cell. **(B)** Western blot of NIH/3T3 cell lines CRISPR-edited with the indicated sgRNAs. **(C)** Representative FISH images of control (pCrispr) NIH/3T3 cells detecting the indicated mRNAs. Blue: cell mask; Red line: outline of nucleus (based on DAPI stain); Yellow: RNA. Scale bar: 10 μm. **(D)** PDI quantifications of *Rab13, Net1, Cyb5r3* and *Ddr2* mRNA distributions from the indicated CRISPR-edited NIH/3T3 cell lines. Only CNBP loss leads to less peripheral RNA distributions. n=20-118. Error bars: SEM. p-values: *<0.05, ****<0.0001, ns: non-significant by Kruskal-Wallis test with Dunn’s multiple comparisons test. **(E)** Representative FISH images of control (pCrispr) MDA-MB-231 cells detecting the indicated mRNAs. Blue: cell mask; Red line: outline of nucleus (based on DAPI stain); Yellow: RNA. Scale bar: 10 μm. **(F)** Western blot of CNBP levels in CRISPR-edited clonal cell lines. **(G)** PDI quantifications of *RAB13, NET1* and *PKP4* mRNA distributions from the indicated CRISPR-edited MDA-MB-231 cell lines. n=38-64. Error bars: SEM. p-values by Kruskal-Wallis test with Dunn’s multiple comparisons test.

To independently confirm these observations, and to rule out the possibility that residual amounts of hnRNP proteins might prevent us from observing any additional functional contributions, we ablated expression of these RBPs using CRISPR-Cas9 genome editing. Guide RNAs (sgRNAs) targeting each candidate RBP were used, and immunoblot analysis verified that this led to undetectable protein levels of the corresponding RBPs (Fig. 2B). We again assessed the distribution of protrusion-localized mRNAs and extended our analysis to include additional transcripts (*Rab13*, *Net1*, *Cyb5r3* and *Ddr2*) (Fig. 2C). Consistent with the results obtained by transient knockdown, loss of hnRNPA2, hnRNPH1 or hnRNPH2 did not affect the distribution of any of the tested mRNAs (Fig. 2D). CNBP loss, on the other hand, significantly reduced the peripheral localization of all tested mRNAs and the effect was consistently observed in two independently isolated clones (Fig. 2D).

To further address whether this is a conserved CNBP role across species, we used CRIPSR to knockout CNBP expression in the human MDA-MB-231 cell line. Two clonal cell populations were isolated, exhibiting undetectable CNBP expression (Fig. 2F), and the distribution of protrusion-targeted mRNAs was assessed (Fig. 2E, G). Again, CNBP loss significantly reduced the peripheral targeting of all tested mRNAs. Additionally, loss of CNBP did not alter the overall *RAB13* and *NET1* mRNA levels or the corresponding protein amounts (Fig. S2), indicating that at least for these transcripts, CNBP plays a specific role in their localization mechanism. Overall, we conclude that both in mouse and human cells, CNBP is important for localization of mRNAs to cytoplasmic protrusions.

### CNBP binds directly to protrusion localized mRNAs through localization sequences in the 3’UTR

The association of CNBP with localization-competent *in vitro* transcribed mRNA fragments suggested that CNBP might exert its role in RNA targeting through directly binding to localization sequences. To address this, we first explored whether we could detect an interaction of CNBP with protrusion-localized mRNAs *in vivo*. To avoid reassociations that can occur during cell lysis, we performed *in vivo* crosslinking with the short-range crosslinker formaldehyde to induce covalent bonds between RNAs and proteins that are in close contact with them. Then, under denaturing conditions, we immunoprecipitated CNBP and measured the amount of co-precipitated protrusion-localized mRNAs using digital droplet PCR (ddPCR). Indeed, several protrusion-localized mRNAs (*RAB13*, *NET1*, *KIF1C*) were enriched in CNBP immunoprecipitates (Fig. 3A). These associations were specific since no binding was observed when immunoprecipitations were carried out with control IgG. Additionally, in the absence of crosslinking, even though similar amounts of CNBP were recovered (Fig. 3A, left panel), no co-precipitated mRNAs were detected (Fig. 3A, right graphs), indicating that under these denaturing conditions interactions were disrupted unless the binding partners were previously covalently linked with formaldehyde due to their close proximity in cells. Therefore, CNBP associates with protrusion-localized mRNAs *in vivo,* likely in a direct manner.

**Figure 3:**
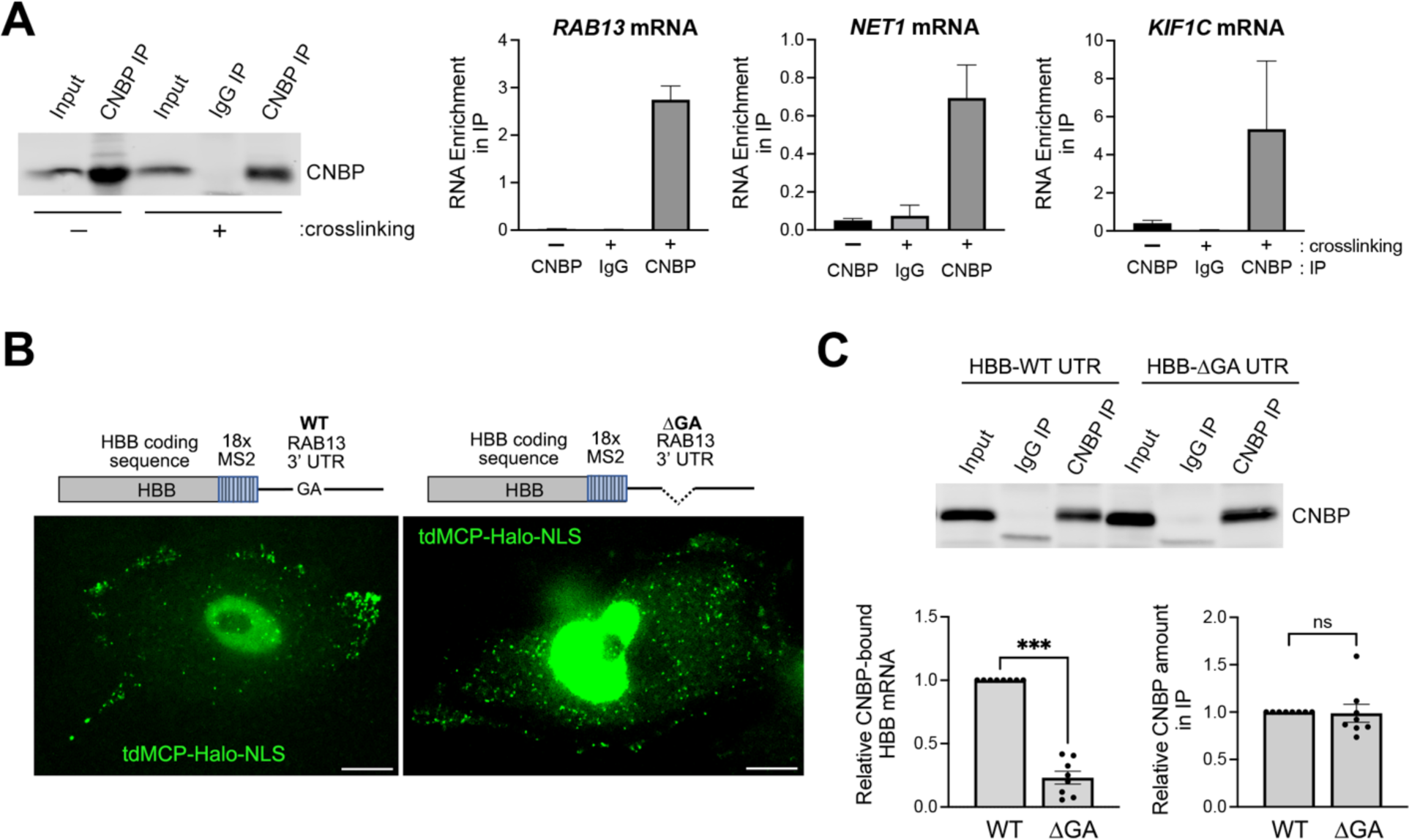
CNBP binds directly to protrusion-localized mRNAs through GA-rich regions. **(A)** MDA-MB-231 cells were crosslinked, or not, with formaldehyde, CNBP was immunoprecipitated and associated RNAs detected by ddPCR. Left panel: Western blot to detect CNBP. Right graphs: amount of indicated mRNAs in immunoprecipitates. Values are expressed as enrichment relative to the corresponding mRNA amount in the input. n=3. Error bars: SEM. **(B)** Schematics of MS2 reporter RNAs containing the Δ-globin (HBB) coding sequence, 18 MS2 hairpins and the WT or truncated RAB13 UTRs. Images are snapshots of cells stably expressing each reporter and tdMCP-Halo-NLS. Cells were labeled with Halo ligand and visualized live. Green spots in the cytoplasm correspond to individual mRNAs. Nuclear signal reflects excess tdMCP-Halo-NLS protein. Scale bar: 10 μm. **(C)** CNBP immunoprecipitation from cells expressing the indicated reporter RNAs. Upper panel: Western blot to detect CNBP. Bottom graphs: amount of reporter mRNA, or CNBP protein, in immunoprecipitates. n=8. Error bars: SEM. p-values: ***<0.001, ns: non-significant by Wilcoxon matched-pairs signed rank test.

To address whether this interaction requires RNA sequences important for protrusion localization, we examined the ability of CNBP to bind to two reporter mRNAs. One contained the full-length 3’UTR of human *RAB13* mRNA while the other contained a truncated UTR (ΔGA) missing the ∼50nt GA-rich sequence that has been shown to be important for peripheral localization ^36^ (Fig. 3B). These reporters also contained the β-globin coding sequence and 18 hairpins of the MS2 bacteriophage for *in vivo* visualization upon co-expression of the MS2 coat protein (MCP) fused to HaloTag (tdMCP-Halo-NLS) (Fig. 3B, schematics). The reporters were stably integrated under an inducible promoter and their expression was induced for a few hours to achieve relatively low levels of expression. Following *in vivo* crosslinking with formaldehyde, CNBP was immunoprecipitated and its association with the reporter RNAs was assessed. As shown in Figure 3C, CNBP bound readily to the reporter carrying the WT *RAB13* 3’UTR but not to the truncated UTR (Fig. 3C, left graph), even though equal levels of CNBP were recovered in each case (Fig. 3C, right graph). Through *in vivo* imaging we verified that the two reporters exhibited distinct distributions in the cytoplasm (Fig. 3B). The reporter carrying the WT 3’UTR accumulated in peripheral protrusions while the one carrying deletion of the GA-rich sequence assumed a more diffuse perinuclear distribution. Importantly, the reduced association of CNBP with the ΔGA-reporter was not due to fact that the mRNA was distributed in the cytoplasm in a way that made it inaccessible to CNBP. In fact, CNBP exhibits a distribution similar to that of the ΔGA-reporter, with the bulk of the protein showing a diffuse perinuclear accumulation by immunofluorescence staining (Fig. S3). Altogether, these data indicate that CNBP binds to 3’UTR localization sequences and is required for peripheral mRNA targeting to cell protrusions.

### CNBP is required for microtubule-dependent mRNA trafficking and associates with KIF1C

mRNA targeting to cell protrusions additionally depends on the KIF1C kinesin motor ^42^. KIF1C co-trafficks with protrusion-targeted mRNAs and is required for their long and directed motions on microtubules ^42^. To determine whether CNBP also affects the microtubule-based transport of protrusion-localized mRNAs, we examined their trafficking upon CNBP knockdown. We have previously reported, using single-molecule mRNA tracking in live cells, that a fraction of mRNAs exhibits long and linear motions over an imaging period of 1 minute in duration. These motions depend on microtubules and KIF1C ^42^. For the studies described here we have constructed an improved MS2-based reporter that incorporates all the regulatory elements of the NET1 transcript: the 5’ and 3’UTRs as well as the NET1A coding sequence (Fig. 4A, schematic). The 3’UTR of NET1 is sufficient to direct protrusion localization of this reporter. Single molecule RNA tracking was performed in cells expressing MCP-Halo upon addition of a fluorescent Halo ligand (Fig. 4A and supplementary movies 1-3). As reported previously ^42^, about 5 percent of tracks per cell were long and directed in control cells and were significantly reduced in cells after transient siRNA-mediated knockdown of KIF1C. Importantly, we found that CNBP knockdown reduced the fraction of long and directed tracks to a similar extent (Fig. 4A and S4). Therefore, both CNBP and KIF1C participate in the microtubule-based trafficking of protrusion-localized mRNAs.

**Figure 4:**
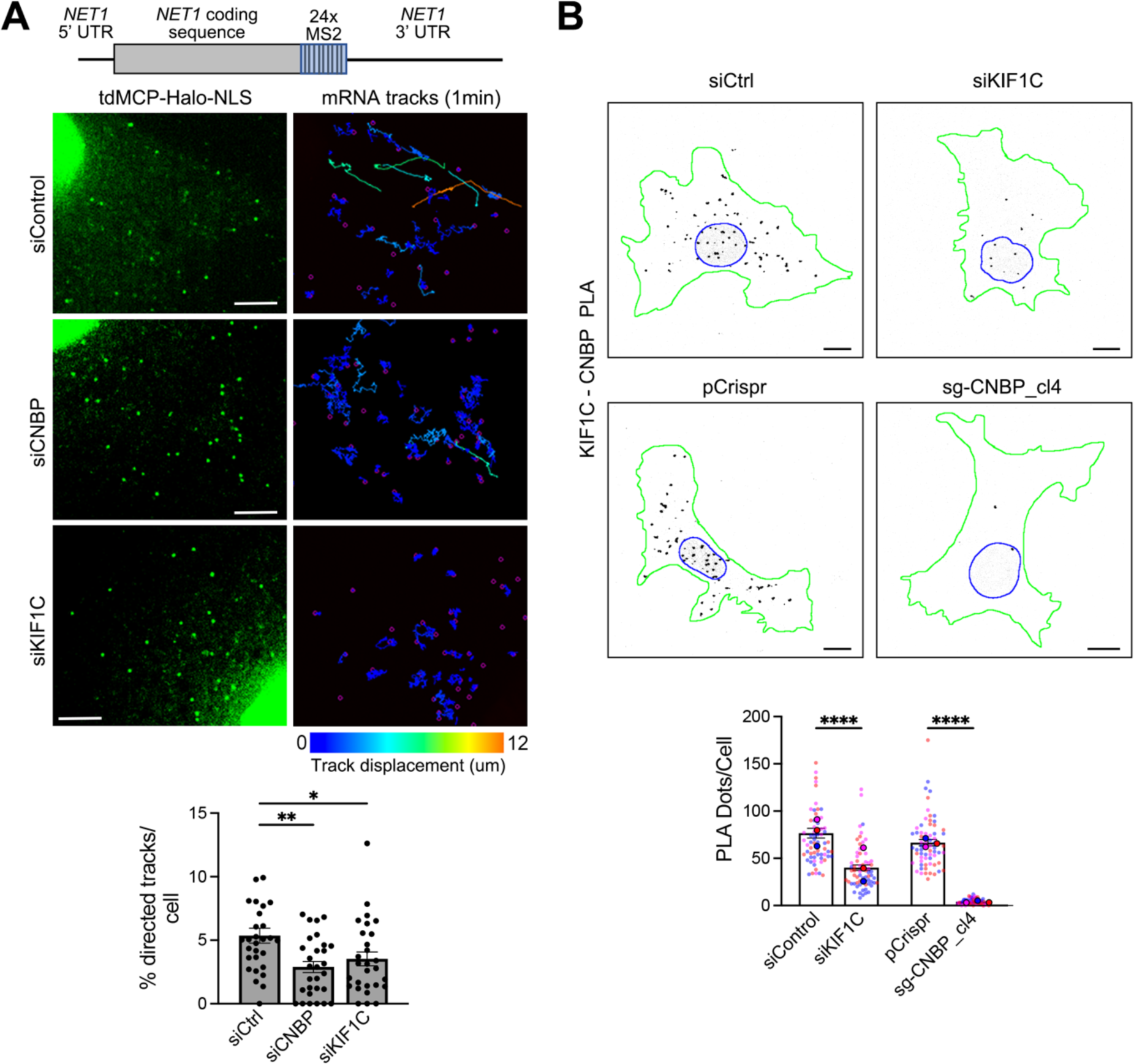
CNBP associates with KIF1C and is required for microtubule-dependent mRNA trafficking. **(A)** Schematic of MS2 reporter RNA containing the NET1 5’- and 3’-UTR and coding sequence as well as 24 MS2 hairpins. Cells stably expressing the reporter and tdMCP-Halo-NLS were transfected with the indicated siRNAs and high-speed imaging was performed over 1 min to track individual RNA movements. Left: images of magnified areas at the beginning of the time lapse. Scale bar: 4 μm. Right: accumulated tracks over the 1 min imaging period. Tracks are color-coded according to total displacement, in μm. Bottom graph: percentage of directed tracks per cell following treatment with the indicated siRNAs. n=28-29. p-values: *<0.05, **<0.01 by Kruskal-Wallis test with Dunn’s multiple comparisons test. **(B)** Representative images of in situ detection of interaction between CNBP and KIF1C by PLA in the indicated CRISPR-edited cell lines, or cells treated with the indicated siRNAs. Black dots: PLA signal; blue outline: nuclear boundary; green outline: cell boundary. Scale bars: 10μm. Bottom graph: quantification of PLA dots per cell. n=65-67 cells in 3 independent experiments. Data points from individual replicates are color coded, and large, outlined color dots indicate the mean of each replicate. Error bars: SEM. p-values: ****<0.0001, by Kruskal-Wallis test with Dunn’s multiple comparisons test.

RBPs can function as adaptors that recruit molecular motors to RNA localization sequences (see Introduction). To gain insights into how CNBP participates in the localization mechanism, we sought to determine whether CNBP and KIF1C physically interact. In co-immunoprecipitation experiments, we could occasionally, but not consistently, detect an interaction of the two proteins (see for example Fig. 5A), suggesting that likely only a small fraction of each of the two proteins are engaged in a complex in cells. We, thus, alternatively interrogated their association by Proximity Ligation Amplification (PLA) assay (Fig. 4B). Using antibodies that recognize the endogenous proteins, we detected substantial PLA signal under control conditions (cells stably expressing only Cas9 (pCrispr) or transiently transfected with a control siRNA (siCtrl)). The observed signal specifically reflected in situ CNBP-KIF1C complexes, since it was significantly reduced in CNBP knockout cells, or upon transient KIF1C knockdown. Interestingly, PLA dots were distributed throughout the cell with a small bias around the nucleus but were noticeably absent from peripheral regions, where protrusion-localized mRNAs accumulate. These results demonstrate that CNBP and KIF1C physically interact in the bulk cytoplasm. They further indicate that their interaction is likely disrupted or altered at peripheral locations.

**Figure 5:**
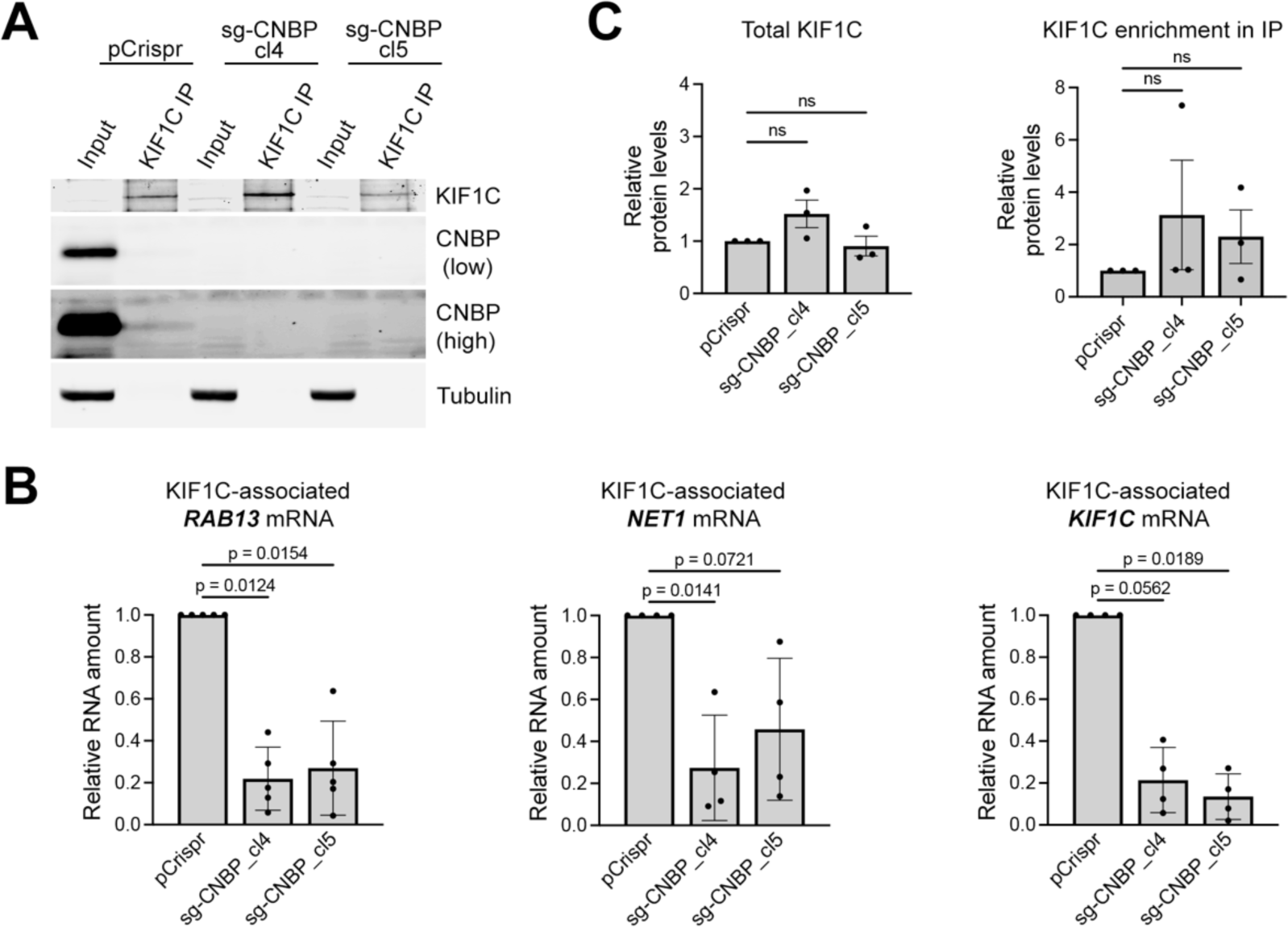
CNBP is required for recruitment of KIF1C to protrusion-localized mRNAs. KIF1C was immunoprecipitated from the indicated CRISPR-edited cell lines (pCrispr control, or CNBP knockout clones). **(A)** Western blot analysis of the indicated proteins from input or IP samples. CNBP blot is shown with adjusted contrast to highlight CNBP protein detected in association with KIF1C (pCrispr-KIF1C IP lane). **(B)** Amount of indicated mRNAs in KIF1C immunoprecipitates by ddPCR. n=4-5. p values by Kruskal-Wallis test with Dunn’s multiple comparisons test. **(C)** Quantification of total KIF1C levels and efficiency of KIF1C immunoprecipitation in control or CNBP knockout cells. Non-significant differences by Kruskal-Wallis test with Dunn’s multiple comparisons test.

### CNBP is required for recruitment of KIF1C to protrusion-localized mRNAs

The fact that CNBP interacts both with RNA localization sequences and with KIF1C suggested that it might function as an adaptor for the recruitment of KIF1C to protrusion-localized mRNAs. To test this idea, we assessed whether loss of CNBP would affect the ability of KIF1C to associate with protrusion-localized mRNAs. For this, we immunoprecipitated KIF1C, either from control cells or the two clonal CNBP knockout cell lines (Fig. 5A) and quantified the number of associated mRNAs by ddPCR (Fig. 5B). Indeed, in the absence of CNBP, KIF1C associated with protrusion-localized mRNAs (*RAB13*, *NET1*, and with its own *KIF1C* mRNA) to a significantly lower degree (Fig. 5B). Importantly, this reduction was not due to changes in the overall expression of KIF1C or the efficiency of KIF1C immunoprecipitation (Fig. 5C). Conversely, loss of KIF1C did not affect binding of CNBP to these mRNAs (Fig. S5). If anything, upon KIF1C loss, CNBP-RNA binding appeared slightly increased. This could potentially reflect that RNAs, which are not trafficked efficiently in the absence of the KIF1C ^42^, remain at perinuclear regions where the bulk of CNBP is also found (see above and Fig. S3) resulting in increased CNBP-RNA association. Regardless of this latter possibility, overall, these data show that CNBP binds to protrusion-localized mRNAs independently of KIF1C but is itself required for KIF1C’s association with protrusion-localized mRNAs, consistent with a role for CNBP as an adaptor for motor recruitment.

### A CNBP-KIF1C motor complex is recruited at GA-rich regions of protrusion-localized mRNAs

The above data suggest that the KIF1C motor is recruited at GA-rich regions of protrusion-localized mRNAs through CNBP. To provide further support for this model, we employed antisense morpholino oligonucleotides (PMOs) that target the GA-rich region of the human *RAB13* mRNA (Fig. 6A). When delivered into cells these antisense oligos disrupt specifically the peripheral localization of the targeted mRNA ^36,37^, however the underlying mechanism of action was unknown. We reasoned that hybridization of these oligos to GA-rich regions might interfere with CNBP-KIF1C complex recruitment. If this prediction is correct, it would support the model of motor recruitment at these UTR regions and would additionally provide a molecular understanding of how antisense oligos interfere with protrusion mRNA localization.

**Figure 6:**
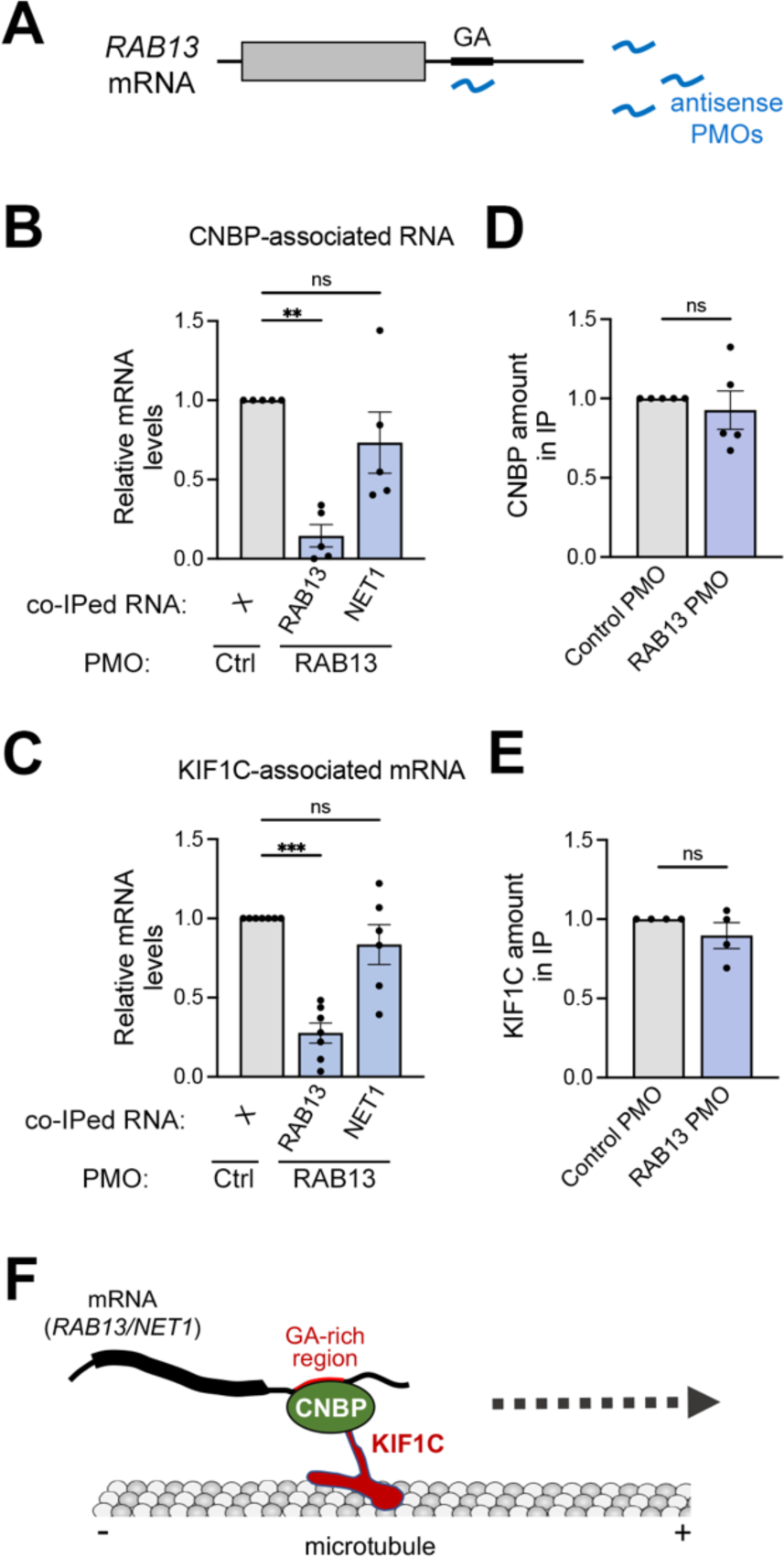
Localization-blocking PMOs against GA-rich regions prevent CNBP and KIF1C binding to target mRNAs. **(A)** Schematic of experimental approach. Antisense PMOs are delivered into cells. PMOs targeting the GA-rich region of the *RAB13* mRNA prevent *RAB13* localization, potentially through interfering with RBP binding. **(B, C)** CNBP (B) or KIF1C (C) immunoprecipitation to detect amount of associated RNAs after PMO delivery. Note that binding to *RAB13* mRNA is specifically affected upon delivery of PMOs targeting the RAB13 GA-region. n=5 (B), 7 (C). Error bars: SEM. p-values: ***<0.001, **<0.01, ns: non-significant by Kruskal-Wallis test with Dunn’s multiple comparisons test. (**D, E)** Relative amount of CNBP (D) or KIF1C (E) protein recovered in immunoprecipitates (IP) by Western blot. ns: non-significant by Wilcoxon matched-pairs signed rank test. **(F)** Proposed model for motor-adaptor complex directing mRNA trafficking to cell protrusions. CNBP binds to GA-rich regions within the 3’UTRs of protrusion-targeted mRNAs and serves as an adaptor for the recruitment of the KIF1C kinesin. KIF1C subsequently traffics mRNAs on microtubules towards the cell periphery.

To address this, either non-targeting control oligos or oligos targeting the GA-rich region of human *RAB13* mRNA were delivered into cells. CNBP or KIF1C proteins were then immunoprecipitated and the number of *RAB13* or *NET1* mRNAs associating with each protein was assessed by ddPCR. Delivery of oligos against *RAB13* led to a significant reduction in the amount of *RAB13* mRNA that associated with CNBP (Fig. 6B), indicating that CNBP binding is prevented by oligo hybridization to the GA-rich region. Importantly, the amount of *RAB13* mRNA that associated with KIF1C was also significantly reduced (Fig. 6C), consistent with the model that KIF1C recruitment relies on CNBP. The observed reduced *RAB13* RNA association with CNBP and KIF1C upon oligo treatment could not be explained by changes in the amount of the corresponding proteins that were immunoprecipitated or expressed in cells (Fig. 6D, E). Furthermore, delivery of oligos targeting the GA-rich region of *RAB13* did not affect binding of CNBP or KIF1C to the *NET1* mRNA, another protrusion-localized mRNA (Fig. 6B, C). These results are consistent with the previously reported specificity of these oligos, which have been shown to affect only their target mRNA ^36–38^. They further indicate that the CNBP-KIF1C motor complex is recruited independently on different mRNAs. Altogether, these results strongly support a model where GA-rich UTR regions provide a platform for the binding of CNBP and the subsequent recruitment of the KIF1C motor for trafficking to cell protrusions (Fig. 6F).

## Discussion

RNA trafficking to mammalian cell protrusions depends on 3’UTR RNA sequences and the KIF1C kinesin ^42^. Here, we have searched for additional components that participate in this transport pathway. We have identified the RNA-binding protein CNBP as a factor that directly associates with localization sequences in the 3’UTR of protrusion-localized mRNAs and mediates their peripheral targeting. Our data indicate that CNBP links individual mRNA cargo to the kinesin motor thus directing their trafficking to cell protrusions. This model is based on the following findings: First, CNBP binds directly to protrusion-localized mRNAs and binding requires the same GA-rich sequences located in their 3’UTR that are needed for proper localization. Second, CNBP interacts with KIF1C both physically and functionally. Third, CNBP is required for KIF1C recruitment to mRNA. Although additional roles of CNBP cannot be excluded, such as regulation of KIF1C motor activity, the evidence presented here supports a simple model of an adaptor-motor complex. This work adds to our understanding of the molecular mechanisms connecting mRNAs to transport machinery for long-range movements in the cytoplasm.

CNBP has been described as a nucleic acid (ssDNA and RNA)-binding protein featuring six to seven tandem CCHC-type zinc knuckle motifs ^45^. An Arg/Gly-rich motif on CNBP is important for binding nucleic acid sequences that are enriched in G nucleotides ^45,46^. Its binding to and unfolding of complex secondary structures, such as G-quadruplexes, has been proposed to facilitate transcription and translation ^45–49^. The work presented here adds on the existing functions of CNBP by describing a novel role in RNA localization by promoting the recruitment of the KIF1C kinesin to RNAs.

We show that CNBP is required for long and linear mRNA movements in the cytoplasm, likely occurring on microtubule tracks. In agreement, we observe that interaction of CNBP with KIF1C is readily observed in the bulk cytoplasmic region, using PLA assay. Interestingly, however, CNBP-KIF1C interaction is not prominently detected in the cell periphery. We have also shown previously that protrusion-targeted mRNAs exist in two physical states: as monomers, that comprise a major fraction of transcript population, and as clusters that are predominantly located at the tips of retracting protrusions and which are heterogeneous, composed of multiple mRNA species ^40^. While KIF1C readily accumulates in peripheral RNA clusters ^42^, we don’t observe a CNBP-KIF1C association in these regions. Altogether, these observations would suggest that when mRNAs reach the cell periphery and become incorporated into KIF1C-containing clusters, CNBP dissociates while KIF1C-RNA association persists.

A few possibilities could underlie such a switch. KIF1C could associate with RNAs through other zinc finger (ZnF)-containing RBPs such as MBNL ^50^. Interestingly, the unstructured carboxy-terminal tail of MBNL supports association with membranes ^50^, raising the intriguing possibility that such a potential RBP switch, from CNBP to MBNL, could maintain KIF1C association while further anchoring the RNA to the proximal plasma membrane at peripheral sites. In a different scenario, KIF1C might directly bind to RNA. Indeed, KIF1C can be crosslinked to mRNAs ^51–53^. Additionally, KIF1C contains a C-terminal intrinsically disorder region (IDR) that drives liquid-liquid phase separation (LLPS) at peripheral protrusive regions ^54^. The purified C-terminal KIF1C IDR can bind and recruit RNA in *in vitro* condensates in a sequence-specific manner ^54^. Thus, the increased local KIF1C concentration at the cell periphery could lead to its condensation and direct association with RNA. In this context it might be relevant that CNBP was recently reported to display antiviral activity against SARS-CoV-2 through binding to the viral RNA and preventing viral RNA and nucleocapsid protein from forming LLPS condensates, which are important for viral replication ^55^. Therefore, one could speculate that CNBP dissociation might be important in allowing KIF1C-RNA condensation at cell protrusions. Delineating how this apparent switch in mode of KIF1C binding is triggered on a molecular level and what functional role it might serve are interesting future questions.

Another aspect of RNA regulation that differs between the bulk cytoplasm and the cell periphery is translation. Protrusion-targeted mRNAs when they are in monomeric form in the bulk cytoplasm are actively translated and exhibit the same translational output independent of their position in the cytoplasm ^40^. On the other hand, peripheral mRNAs found in condensate-like clusters appear translationally repressed as evidenced by their association with fewer nascent polypeptides ^40^. Given that CNBP has been shown to promote translation by binding G-rich elements ^45–47^, we envision that CNBP might additionally participate in translational regulation. In this way, CNBP dissociation from mRNAs at the periphery might lead to translational down-regulation and thus coordinate KIF1C-dependent RNA condensation with translational repression. Therefore, CNBP might have a dual role in the regulation of protrusion-localized mRNAs, by serving as a kinesin adaptor involved in their trafficking and as a spatial regulator of their translation.

Finally, alterations in the CNBP locus have been associated with myotonic dystrophy (dystrophia myotonica, DM) type 2 (DM2) ^56^. Specifically, an expansion of hundreds to thousands of CCTG repeats within intron 1 of the CNBP gene results in intron retention and is thought to be the cause of the condition ^56^. Similarly, a CTG repeat expansion in the 3’UTR of DMPK gene has been linked to myotonic dystrophy type 1 through the production of a toxic transcript that forms foci in the nucleus and sequesters muscleblind-like (MBNL) family of RBPs ^57,58^. In turn, functional compromise of MBNL results in dysregulation of the metabolism of several mRNAs. It has been proposed that a similar indirect, MBNL-dependent mechanism can underlie at least part of the DM2 phenotype. However, a direct role of CNBP cannot be excluded. Although the intronic repeat expansion in CNBP does not alter the amount of transcript produced or its export to the cytoplasm ^59^, reduced levels of protein expression have been reported ^48,60^. Importantly, heterozygous or homozygous CNBP knockout mouse models recapitulate a variety of muscle features observed in DM2 ^61,62^, supporting the idea that at least part of the DM2 defects can be a direct consequence of compromised CNBP protein expression. In light of the roles of both CNBP and MBNL as KIF1C adaptors and the dual role of CNBP as a translation regulator and a kinesin adaptor discussed here, this work provides the basis for interesting hypotheses towards understanding disease mechanisms.

## Acknowledgements

We thank the CCR Genomics Core of the National Cancer Institute, NIH for ddPCR analysis. This work was funded by the Intramural Research Program of the Center for Cancer Research, National Cancer Institute (NCI), National Institutes of Health (NIH) (1ZIA BC011501 to S.M.)

## Author contributions

S.M., K.M. and T.W. designed experiments. K.M., T.W., M.S. and A.N.G. performed and analyzed experiments. L.J. and E.L.P. performed and analyzed mass spectrometry. K.M., S.M. and A.N.G. wrote and edited the manuscript. All authors reviewed the manuscript.

## Supplemental Information

**Table S1:** Excel file containing results of mass spectrometry analysis from two replicate experiments.

**Table S2:** DNA sequences used for generation of in vitro transcription templates

**Movie S1:** Single molecule RNA tracking in siControl-tranfected cells, related to Figure 4A. Cell expressing the NET1 RNA reporter and tdMCP-Halo-NLS imaged at 6.67 frames per second for 1 min. Left panel: raw signal. Right panel: raw signal with overlaid accumulated tracks of individual RNA spots. Only tracks lasting longer than 2.5 seconds are shown. Warmer colors indicate tracks of higher displacement. Scale bar: 5 um.

**Movie S2:** Single molecule RNA tracking in siCNBP-tranfected cells, related to Figure 4A. Cell expressing the NET1 RNA reporter and tdMCP-Halo-NLS imaged at 6.67 frames per second for 1 min. Left panel: raw signal. Right panel: raw signal with overlaid accumulated tracks of individual RNA spots. Only tracks lasting longer than 2.5 seconds are shown. Warmer colors indicate tracks of higher displacement. Scale bar: 5 um.

**Movie S3:** Single molecule RNA tracking in siKIF1C-tranfected cells, related to Figure 4A. Cell expressing the NET1 RNA reporter and tdMCP-Halo-NLS imaged at 6.67 frames per second for 1 min. Left panel: raw signal. Right panel: raw signal with overlaid accumulated tracks of individual RNA spots. Only tracks lasting longer than 2.5 seconds are shown. Warmer colors indicate tracks of higher displacement. Scale bar: 5 um.

## STAR Methods

### RESOURCE AVAILABILITY

#### Lead Contact

Further information and requests for resources and reagents should be directed to and will be fulfilled by the Lead Contact, Stavroula Mili.

#### Materials Availability

All unique/stable reagents generated in this study are available from the Lead Contact with a completed Materials Transfer Agreement.

#### Data and Code Availability

This paper does not report original code.

Any additional information required to reanalyze the data reported in this paper is available from the Lead Contact upon request.

### EXPERIMENTAL MODEL AND SUBJECT DETAILS

#### Cell culture

MDA-MB-231 cells were obtained from ATCC (cat # HTB-26) and cultured in Leibovitz’s L-15 medium (Invitrogen cat# 11415064) supplemented with 10% FBS (or Tet-approved FBS for cells expressing inducible reporter RNAs) and 1% Penicillin-Streptomycin at 37°C in atmospheric air in a humidified cell culture incubator. NIH/3T3 mouse fibroblast cells were cultured in DMEM (Invitrogen cat# 1995073) supplemented with 10% calf serum, sodium pyruvate and 1% Penicillin-Streptomycin at 37°C and 5% CO_2._ Cells were passaged by trypsinization using 0.05% trypsin (Invitrogen cat# 25300120). Cells used in this study have tested negative for mycoplasma.

#### Cell lines

To generate stable cell lines expressing RNA reporters, MDA-MB-231 cells were infected with lentivirus expressing stdMCP-stdHalo (Addgene #104999, modified to remove the Kozak sequence and ATG initiating codon from the sequence of the first HaloTag). A uniformly expressing cell population was selected by fluorescence activated cell sorting. This population was then infected with pInducer20-based constructs expressing Δ-globin followed by 18xMS2 binding sites and either the wild type human RAB13 3’UTR (accession#: NM_002870.5) or a deletion mutant lacking a GA-rich region of the RAB13 3’UTR (corresponding to nucleotides 202-254). NIH/3T3 cells were also infected with stdMCP-stdHalo and with a pInducer20-based construct expressing the NET1A 5’UTR, coding sequence and 3’UTR together with 24xMS2v7 repeats after the coding sequence. Stably expressing lines were selected with Geneticin (Thermo Fisher Scientific). Expression of the reporters was induced by addition of 1 μg/ml doxycycline approximately 3-4 hrs before imaging.

To generate CRISPR edited lines, NIH/3T3 or MDA-MB-231 cells were infected with pLentiCRISPRv2 expressing appropriate guide RNAs and selected with puromycin. Where indicated individual clonal cell lines were isolated by limited dilution.

### METHOD DETAILS

#### Plasmid constructs and lentivirus production

To generate templates for in vitro transcription, inserts containing a T7 promoter, and the indicated UTRs followed by a BoxB sequence were cloned into HindIII and EcoRI sites of pTZ19R (exact sequences used are indicated in Table S2). Plasmids linearized with EcoRI were used as templates for in vitro transcription.

To generate lentiviral plasmids expressing Cas9 together with single guide RNAs, the pLentiCRISPRv2 plasmid was used (Addgene #52961). Target sequences of cloned sgRNAs were as follows: mCNBP-Ex3: TGCCAAGGATTGTGATCTGC; mHNRNPA2B1-Ex3: TCTCTTGCTACAGCACGTTT; mHNRNPH1-Ex3: TCAACAAAAGCCTCGCCACT; mHNRNPH2-aa19: TCCTCGGCTGAGCAGGACCA; mHNRNPH2-aa50: CGTTTCATCTACACCAGAGA (targets both hnRNPH1 and hnRNPH2); hsCNBP-Ex2-1: GTGTGGACGATCTGGCCACT. sgRNA sequences were cloned onto BsmBI sites of pLentiCRISPRv2 and plasmids were used for lentivirus production.

Lentiviruses were produced in HEK293T cells. The cells were transfected with the lentiviral vectors together with the pMD2.G and psPAX2 packaging plasmids using PolyJet In Vitro DNA Transfection Reagent (SignaGen) for 48 hrs. Harvested virus was precipitated with polyethylene glycol overnight at 4°C.

#### In vitro transcription and λN-GST pulldown

In vitro transcriptions were carried out from linearized plasmid templates (4 ugr in 50 ul) using T7 Polymerase HC (Promega, cat# P2075) in transcription optimized buffer (Promega) with 20mM DTT, 10mM MgCl_2_, 4mM NTPs, 40U RNasin Plus (Promega, cat# N2615) and 0.1% Triton X-100. The reaction was incubated at 37°C for 3.5hrs.

For λN-GST pulldown, in vitro transcription reactions were incubated with 45ug purified λN-GST protein for 15 min at room temperature. The mix was incubated with 15uL Glutathione magnetic beads (Invitrogen, cat #78602) for 20 min at room temperature with shaking, and the beads and bound RNA were washed with lysis buffer (50mM Tris-Cl pH 7.5, 0.5% Triton-X 100, 100mM NaCl, 2.5mM MgCl_2_) and further blocked with lysis buffer containing E. coli tRNA, BSA (Sigma, cat#10711454001), salmon sperm DNA (Invitrogen, cat# 15632011) and glycogen (Invitrogen, cat# AM9510) for 30-60 mins at 4°C. Cell extracts from MDA-MB-231 or NIH/3T3 cells were prepared by lysing in lysis buffer with Halt protease inhibitor (Invitrogen, cat #78444) and RNase inhibitor (Promega, cat# N2615), brief sonication and clearing at 10,000xg for 10 min at 4°C. The GSH beads and bound RNA were incubated with extract from 5-10×10^6^ cells at 4°C for 1 hour with rotation and washed first with lysis buffer and then with PBS pH 8.0. Bound material was eluted in 50ul of 40mM reduced L-Glutathione in PBS pH 8.0. Bound RNA was analyzed on Agilent Tapestation, and bound protein by Silver staining, Western blot or mass spectrometry.

#### Immunoprecipitation

For KIF1C immunoprecipitation, cells were lysed with a buffer containing 10mM Tris-Cl pH 7.4, 100mM NaCl, 2.5mM MgCl_2_, 0.5% Triton X-100, Halt protease and phosphatase inhibitor cocktail (Invitrogen, cat #78444) and RNase inhibitor (Promega, cat# N2615). Lysates were sonicated (setting 2, 2x 5sec, Misonix Inc. Sonicator XL), cleared by centrifugation at 14,000xg for 10 min at 4°C, and mixed with Protein G Dynabeads (Invitrogen, cat #100004D) pre-bound with 1ug anti-KIF1C antibody (Bethyl cat# A301-070A) and pre-blocked with E. coli tRNA, salmon sperm DNA (Invitrogen, cat# 15632011), BSA (Sigma, cat#10711454001), and glycogen (Invitrogen, cat# AM9510). Lysate and bead mix was incubated for 1.5 hrs at 4°C. Bound material was eluted with lysis buffer containing 1% SDS and processed for protein or RNA analysis.

For CNBP immunoprecipitation, cells were crosslinked with 0.3% paraformaldehyde (Electron Microscopy Sciences, cat #15710) in PBS for 10min at room temperature. Crosslinking was stopped by addition of 250mM glycine pH 7.0, and cells were lysed in RIPA buffer (50mM Tris-Cl pH 7.4, 150mM NaCl, 1% Triton X-100, 0.5% Na-deoxycholate, 0.1% SDS) with Halt protease and phosphatase inhibitor cocktail (Invitrogen, cat #78444) and RNase inhibitor (Promega, cat# N2615). Lysates were sonicated (setting 4.5, 5x 15sec, Misonix Inc. Sonicator XL), cleared by centrifugation at 14,000xg for 10 min at 4°C, and mixed with Protein G Dynabeads pre-bound with 2ug anti-CNBP antibody (Proteintech cat# 67109) and pre-blocked with E. coli tRNA, salmon sperm DNA, BSA, and glycogen. Lysate and bead mix was incubated for 3 hrs at 4°C. Bound material was eluted with a buffer containing 100mM Tris-Cl pH 6.8, 5mM EDTA, 10mM DTT, 1% SDS, incubated at 70°C for 50 min, and processed for protein analysis. For RNA analysis samples were further digested with proteinase K at 37°C for 30 min before Trizol extraction.

#### siRNA and morpholino transfection

For knockdown experiments, 40 pmoles of siRNA were transfected into cells by Lipofectamine RNAiMAX (ThermoFisher, cat# 13778-150) according to the manufacturer’s protocol. Cells were assayed 72 hours post-transfection. The following siRNAs were used: AllStars negative control (Qiagen cat# 1027281), si-Mm-Cnbp #6 (Qiagen cat# SI02672313; target sequence: 5’- CAGCAAGACAAGTGAAGTCAA -3’), si-Mm-hnRNPA2B1 #3 (Qiagen cat# SI00210672; target sequence: 5’- AAGGCATTGTCTAGACAAGAA -3’), si-Mm-Hnrnph1 #4 (Qiagen cat# SI01068403; target sequence: 5’- TAGGAGCTGCGTCTACAATTA -3’), si-Mm-hnrnph2 #1 (Qiagen cat# SI01068452; target sequence: 5’- ATGTTGTAGGAGTGTACTTAA -3’)

Antisense morpholino oligonucleotides were synthesized by GeneTools, LLC and delivered into cells using EndoPorter(PEG) (GeneTools, LLC). A combination of two oligos were used for RAB13 and NET1, at a final concentration of 20 μM. Sequences used are as follows: Control: 5’-CCTCTTACCTCAGTTACAATTTATA-3’; RAB13: 5’-TCTTTCACTTCCTCAATTCATTCCT-3’ and 5’-CCTTCCTTTCCTCCTCCCTCTCTTC-3’; NET1: 5’-TCCCTCTTGCATTTCAGACAACACT-3’ and 5’-GACAAAACTACTCTCTTTTCCTCTC-3’. Cells were assayed 72 hours post-transfection.

#### Western blot

The following primary antibodies were used: rabbit polyclonal anti-hnRNPH1 (Bethyl, cat# A300-511A; 1:10,000), rabbit monoclonal anti-hnRNPH2 (Abcam, cat# ab179439; 1:1,000), rabbit polyclonal hnRNPF (Abcam, cat# ab50982; 1:1,000), mouse monoclonal hnRNPA2 (Santa Cruz, cat# sc-53531; 1:1,000), mouse monoclonal anti-CNBP (Proteintech, cat# 67109; 1:3000), rabbit polyclonal anti-CNBP (ThermoFisher, cat# PA5-35241; 1:1,000), rabbit polyclonal anti-KIF1C (Bethyl, cat #A301-070A; 1:1000), mouse monoclonal hnRNPK (Santa Cruz, cat# sc-28380), rabbit monoclonal anti-GAPDH (Proteintech, cat# 2118). Anti-rabbit and anti-mouse secondary antibodies from Li-Cor were used at 1:10,000. Membranes were scanned using an Odyssey fluorescent scanner (Li-Cor) and bands were quantified using ImageStudioLite (Li-Cor).

#### Mass spectrometry

Proteins were denatured by addition of 8M urea in 100 mM ammonium bicarbonate and then heating at 37 °C for 15 min. They were then reduced by reaction with 10 mM DTT for 1 h at 37 °C and reduced by reaction with 50 mM iodoacetamide for 30 min at room temperature. The urea concentration was reduced to 1 M and proteins trypsin digested overnight at 37 °C. Following digestion, the peptides were desalted on a C18 spin desalting column and dried by lyophilization. Dried peptides were resuspended in 5% acetonitrile, 0.05% TFA in water for mass spectrometry analysis on an Obitrap Fusion Tribrid (Thermo) mass spectrometer. The peptides were separated on a 75 µm x 15 cm, 3 µm Acclaim PepMap reverse phase column (Thermo) at 300 nL/min using an UltiMate 3000 RSLCnano HPLC (Thermo) and eluted directly into the mass spectrometer. For analysis, parent full-scan mass spectra collected in the Orbitrap mass analyzer set to acquire data at 120,000 FWHM resolution and HCD fragment ions detected in the ion trap. Proteome Discoverer 2.0 (Thermo) was used to search the data against the murine database from Uniprot using SequestHT. The search was limited to tryptic peptides, with maximally two missed cleavages allowed. Cysteine carbamidomethylation was set as a fixed modification, with methionine oxidation as a variable modification. The precursor mass tolerance was 10 ppm, and the fragment mass tolerance was 0.8 Da. The Percolator node was used to score and rank peptide matches using a 1% false discovery rate.

#### RNA isolation and ddPCR

RNA was extracted using Trizol LS reagent (Invitrogen, cat # 10296028) according to the manufacturer’s instructions. Prior to extraction an equal amount of exogenous spike RNA (in vitro transcribed beta-globin or GFP RNA) was added to each sample to correct for differences in recovery during the process. Isolated RNA was treated with RQ1 DNAse (Promega, cat #M6101) for 30min at 37°C and purified again with Trizol. RNA was reverse transcribed using the iScript cDNA Synthesis Kit (Bio-Rad, cat# 1708891). For ddPCR, cDNA samples were analyzed using the ddPCR EvaGreen Supermix (Bio-Rad, cat. no. 186-4034). Droplets were generated using the Automated Droplet Generator (Bio-Rad, cat no. 186-4101), PCR amplification was performed on a C1000 Touch™ Thermal Cycler (Bio-Rad, cat no. 185-1197) and droplet reading was done with the QX 200 Droplet reader (Bio-Rad, cat no. 186-4003) and QuantaSoft software (Bio-rad). Specificity of primers pairs in detecting RAB13, NET1, KIF1C, beta-globin and GFP RNAs was verified by comparing with RNA from knockdown cells.

#### RNA Fluorescence in situ hybridization (FISH)

MDA-MB-231 or NIH/3T3 cells were plated on either collagen IV (Sigma, cat# C5533; 10ug/mL) or fibronectin (Sigma, cat# F1141; 5ug/mL) coated coverslips, respectively, for 2hrs and fixed with 4% paraformaldehyde for 20 mins at RT. FISH was performed using the ViewRNA ISH Cell Assay kit (ThermoFisher, cat# QVC0001) according to the manufacturer’s protocol. The following probes were used in this study: mouse Rab13 (cat# VB1-14374-01), mouse Net1 (cat# VB1-3034209-01), mouse Cyb5r3 (cat# VB1-18647-01), mouse Ddr2 (cat# VB1-14375-01), human RAB13 (cat# VA1-12225-06), human NET1 (cat# VA1-20646-01) and human PKP4 (cat# VA1-12406-01). HCS Green cell mask (Invitrogen, cat# H32714) was used to identify the cell border and samples were mounted in ProLong Gold antifade with DAPI (Invitrogen, cat# P36931).

#### Proximity Ligation Assay (PLA)

MDA-MB-231 cells plated on collagen IV-coated (10ug/mL) coverslips were washed with PBS and fixed for 15 min at RT with 4% paraformaldehyde then permeabilized for 5 min at RT with 0.2% Triton X-100. The DuoLink In Situ Red Kit (Sigma, cat #DUO92008) was used for PLA and the manufacturer’s protocol was followed. Briefly, the cells were blocked using the provided blocking buffer at 37°C for 1hr in a humidified chamber. Primary antibodies were diluted in the provided DuoLink antibody diluent and incubated on the cells for 1.5hrs at RT in a humidified chamber. The following primary antibodies were used: rabbit anti-KIF1C (Proteintech, cat# 12760-1-AP; 1:100), mouse anti-CNBP (Proteintech, cat# 67109-1-IG; 1:500). After washing, the PLA probes supplied with the kit were used at a 1:10 dilution in the antibody diluent and incubated for 1hr at 37°C. Ligation was performed for 30 min at 37°C then amplification was performed for 100 min at 37°C. The cells were washed and fixed again in 4% paraformaldehyde in PBS for 10 min at RT then stained with Alexa Fluor 488 Phalloidin (Invitrogen, cat# A12379; 1:500) in blocking buffer for 20 min at RT. Coverslips were mounted in the provided DuoLink mounting medium with DAPI and kept at 4°C in the dark until imaging the next day.

#### Microscopy and Image Analysis

RNA FISH and PLA experiments were imaged on a Leica SP8 confocal microscope with an HC PL APO 63x oil immersion objective. Z-stacks through the cell volume were obtained and maximum intensity projections were used for all analysis. For FISH images, calculation of the PDI index was performed using a previously published custom Matlab script^44^. For PLA, an ImageJ script based on the Analyze Particles plugin was used to calculate the number of PLA dots present within cells.

Live imaging of cells expressing single-molecule RNA reporters was done using a Nikon Eclipse Ti2-E inverted microscope, equipped with a motorized stage, a Yokogawa CSU-X1 spinning disk confocal scanner unit, and operated using NIS-elements software. Acquisitions were performed using an Apochromat TIRF 100× oil immersion objective (N.A. 1.49, W.D. 0.12 mm, F.O.V. 22 mm) and Hamamatsu ORCA-Fusion BT Gen III back-illuminated sCMOS cameras. Constant 37°C temperature and 5% CO_2_ were maintained using a Tokai Hit incubation system. To label MCP-Halo proteins, cells were supplemented with 200nM of JFX554 HaloTag ligand, obtained from Janelia Research Campus for 2 hrs. The medium was then replaced, and 1 mg/ml doxycycline was added to induce expression of reporter mRNAs for 3 hrs. Cells were plated on fibronectin (5 mg/mL)-coated 35 mm glass bottom dishes for ∼1 hr, and samples were excited using a 488 nm (20mw) laser line and imaged at a rate of 6.66 fps for 60sec.

Single molecule RNA tracking was performed using TrackMate plugin in ImageJ/Fiji. For every cell, all tracks lasting for >2.5 secs (ca. 17 consecutive frames) were used for analysis. Values of ‘Track displacement’, ‘Linearity of forward progression’ and ‘track duration’ were extracted and plotted. ‘Track displacement’ is defined as the distance from the first to the last spot of the track. ‘Linearity of forward progression’ is the mean straight line speed divided by the mean speed; where mean straight line speed is defined as the net displacement divided by the total track time.

#### Statistical analysis

All statistical analysis was performed using GraphPad Prism software using the appropriate statistical tests as indicated within the text and figure legends. Normally distributed datasets were analyzed using parametric statistical tests. Datasets deviating from a normal distribution were analyzed using non-parametric tests. Follow up tests were included, as appropriate, to adjust for multiple comparisons.

## Supplementary Figures and Legends

**Figure S1:**
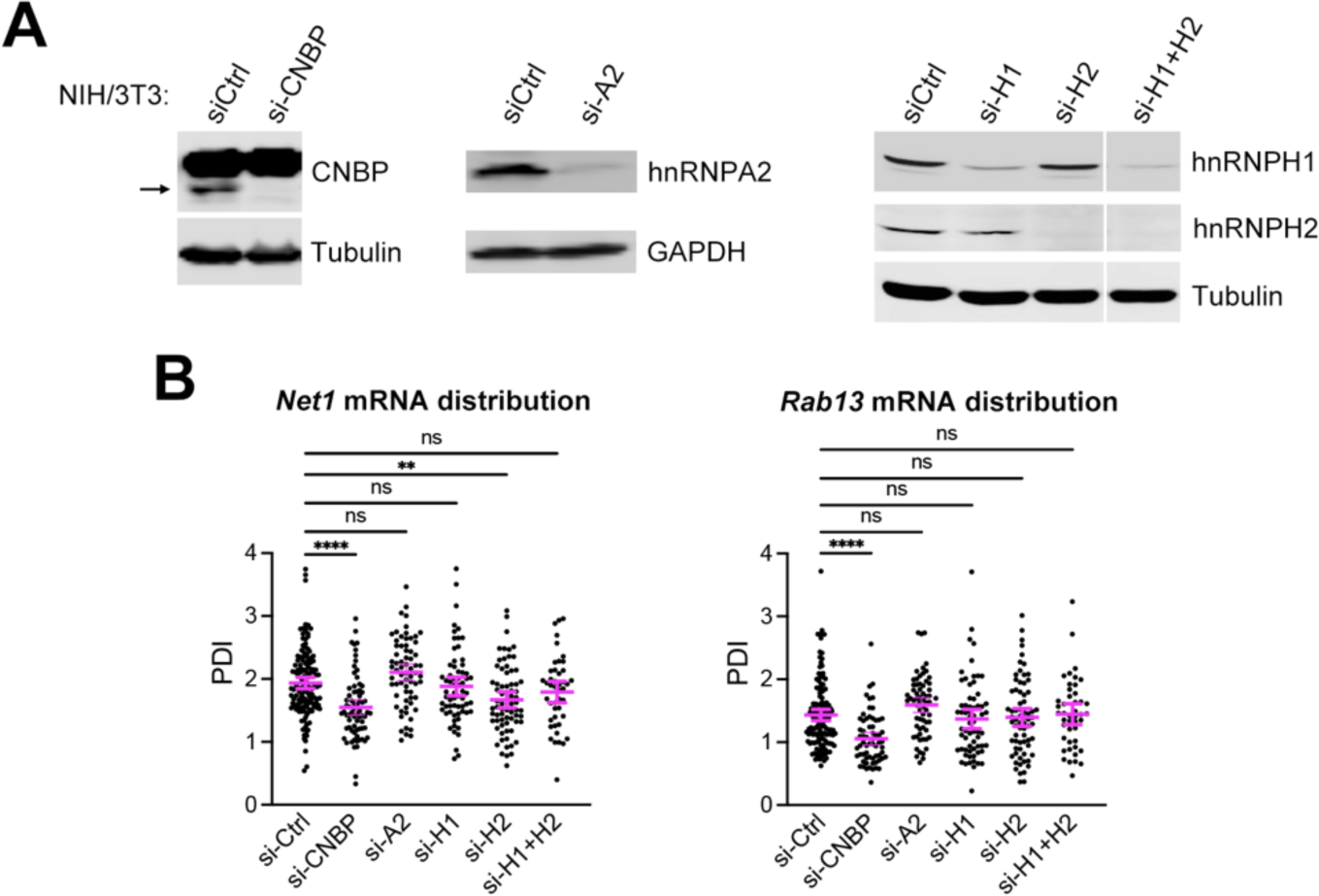
CNBP knockdown prevents localization of mRNAs to cytoplasmic protrusions. **(A)** Western blot of NIH/3T3 cells transfected with the indicated siRNAs against RBPs (CNBP, hnRNPA2, hnRNPH1 and hnRNPH2). Note that the CNBP antibody used cross-reacts prominently with a non-specific band. Signal corresponding to CNBP is indicated by an arrow. **(B)** PDI quantifications of *Rab13* and *Net1* mRNA distributions from the indicated siRNA-treated cells. n=45-136. Error bars: SEM. p-values: **<0.01, ****<0.0001, ns: non-significant by Kruskal-Wallis test with Dunn’s multiple comparisons test.

**Figure S2:**
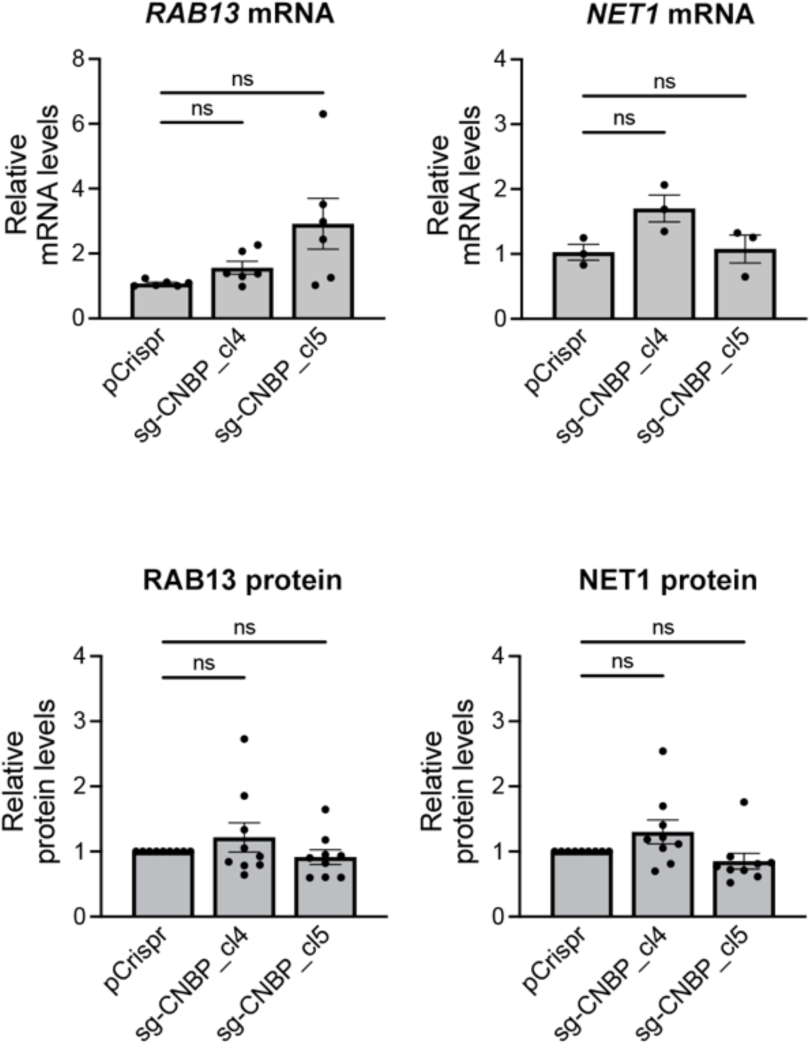
CNBP loss does not affect RAB13 and NET1 mRNA or protein levels. RAB13 and NET1 mRNA levels (measured by ddPCR) and protein levels (measured by Western blot) in the indicated CRISPR-edited clonal cell lines.

**Figure S3:**
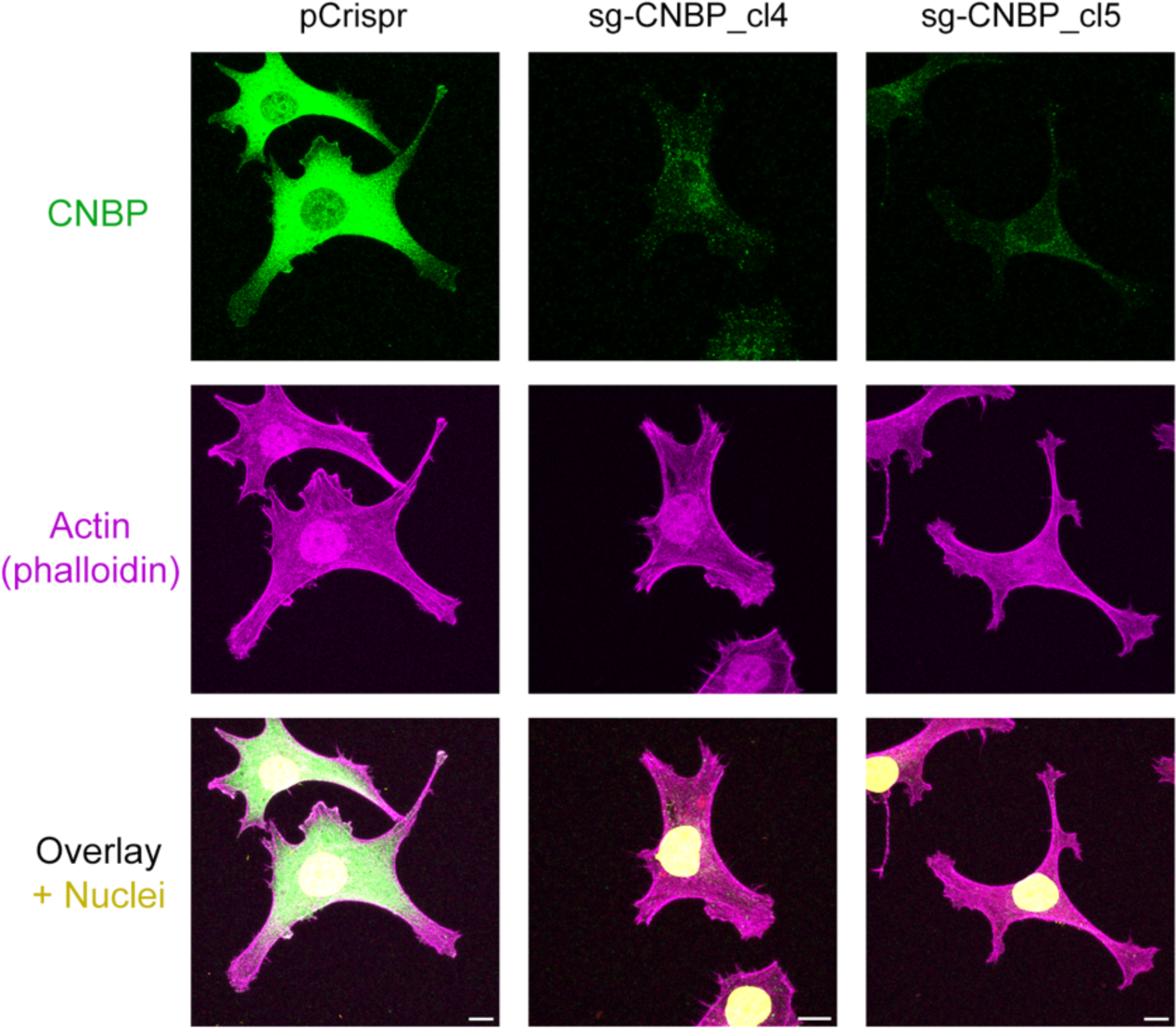
CNBP distributes diffusely in the perinuclear cytoplasm. CNBP immunofluorescence in the indicated CRISPR-edited clonal cell lines. Cells were also stained with phalloidin to visualize the actin cytoskeleton and delineate cell morphology, as well as with DAPI to visualize nuclei. Staining is specific for CNBP since it is markedly reduced in CNBP knockout clones. CNBP distributes diffusely in the cytoplasm with the bulk accumulating around the nucleus. Scale bars: 10μm.

**Figure S4:**
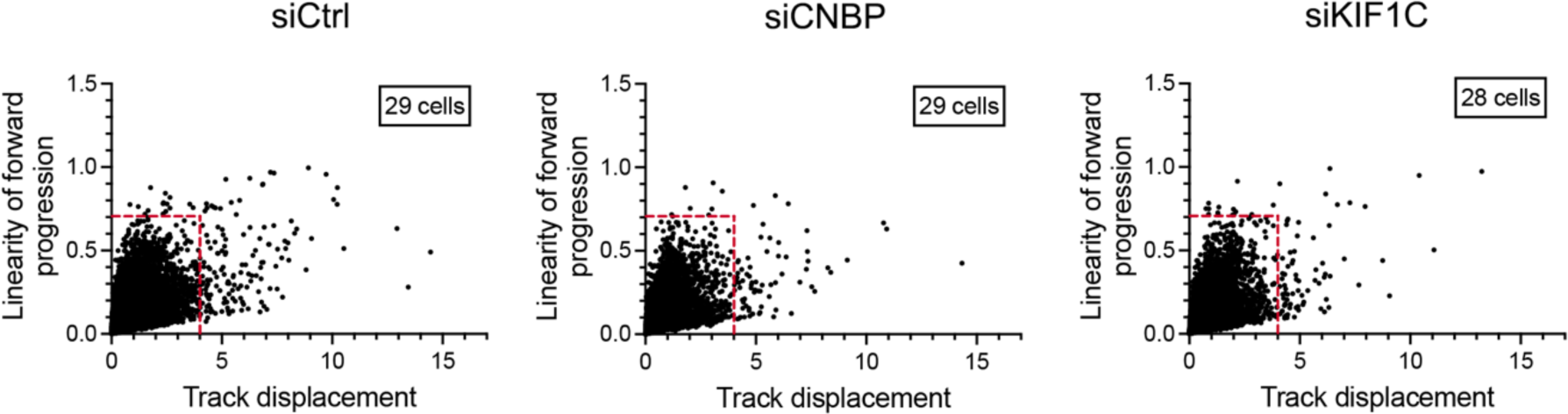
RNA particle tracking metrics. Graphs plot the displacements of all individual RNA tracks (*x*-axis) over the linearity of their forward progression (*y*-axis) (defined as the mean straight line speed divided by the mean speed), from cells expressing MS2 RNA reporter and treated with the indicated siRNAs. Representative images and particle tracking movies are shown in Fig. 4A and supplementary movies 1-3. Red lines indicate the thresholds used to filter tracks of molecules undergoing directed movement (based on ref ^42^).

**Figure S5:**
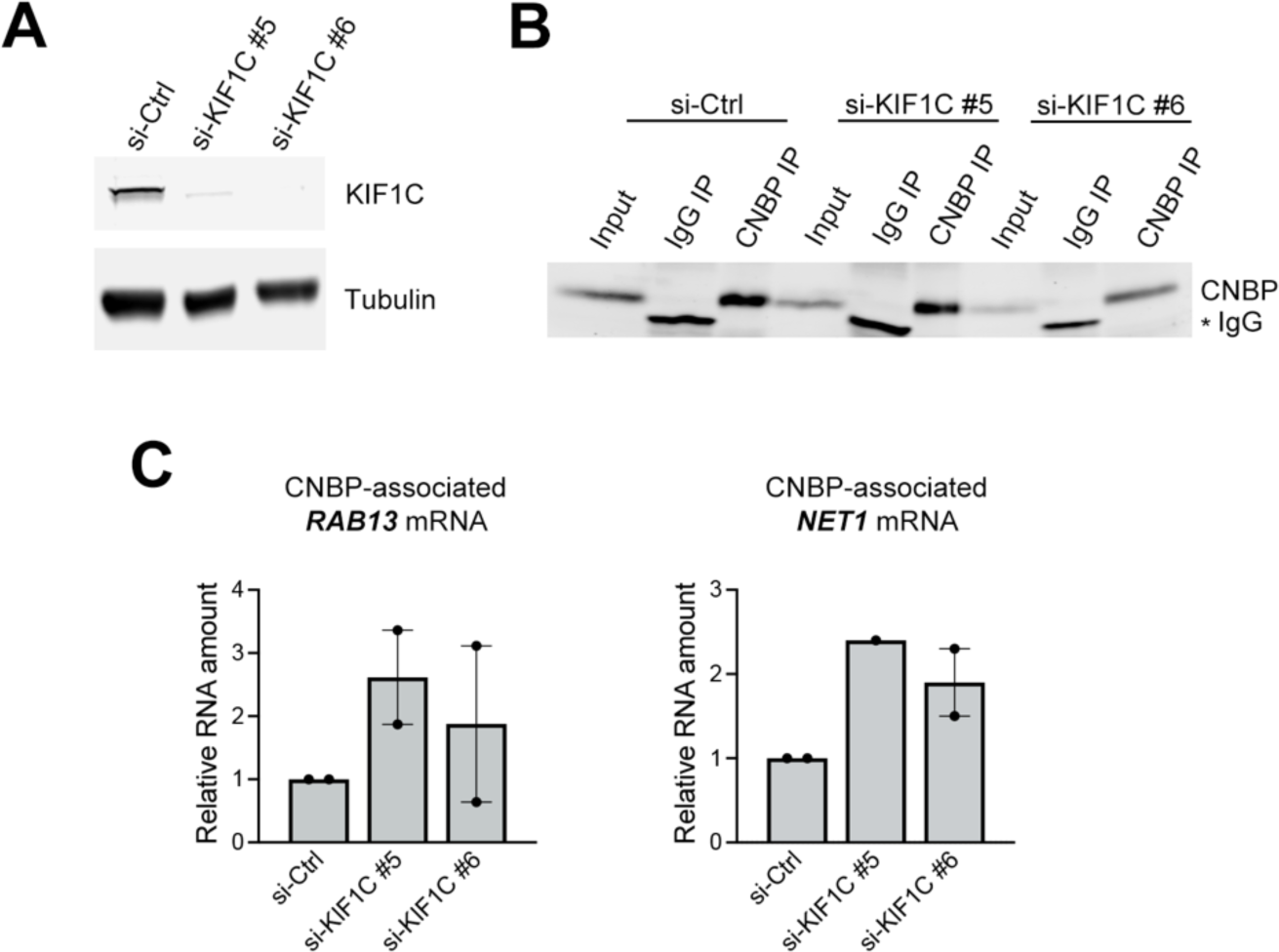
KIF1C is not required for binding of CNBP to protrusion-targeted mRNAs. **(A)** Western blot of MDA-MB-231 cells transfected with the indicated siRNAs. **(B)** Control or si-KIF1C transfected cells were immunoprecipitated with anti-CNBP, or control IgG, antibodies and recovered protein was analyzed by Western blot. Asterisk indicates a band originating from the IgG antibody. **(C)** Amount of *RAB13* and *NET1* mRNAs co-immunoprecipitated with CNBP from cells treated with the indicated siRNAs. Measured by ddPCR and expressed as amount of RNA in IP eluate over amount in the input normalized to control sample.

