## Supplementary material for "A KIF1C-CNBP motor-adaptor complex for trafficking mRNAs to cell protrusions": Table S2

**Table S2:** Sequences of inserts used for generation of in vitro transcription templates.

**T7prom-MmPkp4-BoxB**

**AAGCTTTAATACGACTCACTATAG**CATCAAGACGGCTGCCTGCTGAGGGGCGCTTTCCTTCT
GACTCTGTTTGGATTGAGGGGAAGTCCGTCTTGCTGATGATGGTGACCGTGAAAGTGAAATGG
AAGGGATGAGTGAAGAGGTTTTGGTTTGTTTGTTTGTTTTTTCTTTTTTTGAGGAATTTTCAGGGAA
GTGAGGAAACCCTTGGGAGAGGACTTTGTACGCGCTGTGTAGGTGTTAGATCTAATTACTTGTA
GAGTCTAGTGGTGAAGGTGTGGGTGACGTGCTGGGAGGCTTGAGACGTGGGTGAGATGAGAT
GGGTATGTGTAGGTCAAATCAAATGACAGATGATTTTTTTAATGTGAATAAAGTTATGTTCAGATA
GTTTGTACAGAAAAAATAATAATAAAATGGATGCCCTTCATGTTTTATTGCTATTACTAAATGTCAA
GATTGTATGCTATTATGTCTTGTAAAAATTCCTTCTGTTGGTGTAAATATGGAAATGCCACATTGGT
TAAGTGCCATCAATTGTAATGCAGTGTGTCATTTGAAAAGAGATTTGAAGAACTGACAGCTTAAA
GCCCAAGCGGGAAACCCGCCCGGGAACTGTTCGCAGTTGACAACAAACTCTGACGCCCCTC
TGTTTTCAGTGAGTAGTGAACTCCGGAAGCACAAAGGCCAGCGTGACAGCAGCGCCATGCT
CATCCCCCTCACAGGACACTTCACTGCCATTTTCTATGCACATGGAAGAATAATAAATGTGGAA
ATTTCATCCTGAA**GCGCCGAACT**GGGCCCgacgactgtagaaaagggccctgaagaagggccctctgctgtct
agcGAATTC

**T7prom-MmPkpA-BoxB**

**AAGCTTTAATACGACTCACTATAG**CATCAAGACGGCTGCCTGCTGAGGGGCGCTTTCCTTCT
GACTCTGTTTGGATTGAGGGGAAGTCCGTCTTGCTGATGATGGTGACCGTGAAAGTGAAATGG
AAGGGATGAGTGAAGAGGTTTTGGTTTGTTTGTTTGTTTTTTCTTTTTTTGAGGAATTTTCAGGGAA
GTGAGGAAACCCTTGGGAGAGGACTTTGTACGCGCTGTGTAGGTGTTAGATCTAATTACTTGTA
GAGTCTAGTGGTGAAGGTGTGGGTGACGTGCTGGGAGGCTTGAGACGTGGGTGAGATGAGAT
GGGAA**GCGCCGAACT**GGGCCCgacgactgtagaaaagggccctgaagaagggccctctgctgtctagcGAATTC

**T7prom-MmPkpB-BoxB**

**AAGCTTTAATACGACTCACTATAG**CCCTTCATGTTTTATTGCTATTACTAAATGTCAAGATTGTAT
GCTATTATGTCTTGTAAAAATTCCTTCTGTTGGTGTAAATATGGAAATGCCACATTGGTTAAGTGC
CATCAATTGTAATGCAGTGTGTCATTTGAAAAGAGATTTGAAGAACTGACAGCTTAAAGCCCAA
GCGGGAAACCCGCCCGGGAACTGTTCGCAGTTGACAACAAACTCTGACGCCCCTCTGTTTTCA
GTGAGTAGTGAACTCCGGAAGCACAAAGGCCAGCGTGACAGCAGCGCCATGCTCATCCCC
CTCACAGGACACTTCACTGCCATTTTCTATGCACATGGAAGAATAATAAATGTGGAAATTTCATC
CTGAA**GCGCCGAACT**GGGCCCgacgactgtagaaaagggccctgaagaagggccctctgctgtctagcGAATTC

**T7prom-MmRab13-BoxB**

**AAGCTTTAATACGACTCACTATA**GAGCATTTCTTGCCTCCTATTCACCCTGAACCTGGAGGCT
AGACCTGAGGGAGTCGGACTGAGGGATTGCAGATGGGAGAACTGTGGTGGCACCTCAAGGG
GAGATGAGGGGAATAAGGAGACCGGCGAGGACGAGACGGAAGAAAGGGGCAGGGAAAGG
AGGGGGAGGAACCAAGGATGTGAAAGGTGAACAGAAGGGATTTGAGAAGAGGAAAGGAAGA
AGAAATGAATGGCTCAGGCCTTGGACAGTCCAACATTAAAGTCAACATGCTGATCTCTCCATT
CCTGGTTCAGGGTTAGGGTCCTGAGAGGCTGGCTCGGCACTACTCCGAGGGTCCCTCACTC
TACAAGGTCTTTGTTAGTATTAAAGGCCACTGTTTTGCATGAATGTCCCATTTGCATTACTTTCATT
ATTGTCAGAATTGCTCTTCACTCAAATCCTATTTTTGTCACGCCAAGATATTGGTTCACCTGAATG
TGGCTGGGTTCCCCTTCCTTGCCCCAACTCTTTCACTGGTGATGAAAACAGCATGGGGCAGC
CTGAAGGACGGACATCCTGTTTCCACTGTGGGTTCCCAAGGACTACAAGAGTGGACGGAAC
CTTGCCTTGAGCACACAGTAACCCAAGGACAAAGGATTTGAACCAGGCTTCAGTAAACAGCA
GCACTTAGTATGGTTTATCCAAGGAGATGTGGGACATCTTTGATTCTGATGTAGTCAGCTTAGGT
GTTGGGTACTGTTAGCTGCTTTTGTTAGAGTATTCTCAGTGTTGCACAAAGAAATACATGAACAA
GGTGAA**GCGCCGAACT**GGGCCCgacgactgtagaaaagggccctgaagaagggccctctgctgtctagcGAATT
C

**T7prom-HsRAB13-BoxB**

aagctt**TAATACGACTCACTATA**GGGACCCTTTCTTGCCTCCCCACCCCGGAAGCTGAACCTG
AGGGAGACAACGGCAGAGGGAGTGAGCAGGGGAGAAATAGCAGAGGGGCTTGGAGGGTCA
CATAGGTAGATGGTAAAGAGAATGAGGAGAAAAAGGAGAAAAGGGAAAAGCAGAAAGGAAAA
AAAGGAAGAGAGAGGAAGGGAGAAGGGAGAGGAATGAATTGAGGAAGTGAAAGAAGGCAAG
GAGGTAGGAAGAGAGGGAGGAGGAAAGGAAGGAGAGATGCCTCAGGCTTCAGACCTTACCT
GGGTTTTCAGGGCAAACATAAATGTAAATACACTGATTTATTCTGTTACTAGATCAGGTTTTAGGG
TCCTGCAAAAGGCTAGCTCGGCACTACACTAGGGAATTTGCTCCTGTTCTGTCACTTGTCATG
GTCTTTCTTGGTATTAAAGGCCACCATTTGCACAAATGTTCCTGTTTTGGGTAACTTGGATTATTG
TCAGGGGCCCgacgactgtagaaaagggccctgaagaagggccctctgctgtctagcGCTAGCGCGGCCGC
gaattc

**T7prom-HsRAB13(ΔGA)-BoxB**

aagctt**TAATACGACTCACTATA**GGGACCCTTTCTTGCCTCCCCACCCCGGAAGCTGAACCTG
AGGGAGACAACGGCAGAGGGAGTGAGCAGGGGAGAAATAGCAGAGGGGCTTGGAGGGTCA
CATAGGTAGATGGTAAAGAGAATGAGGAGAAAAAGGAGAAAAGGGAAAAGCAGAAAGGAAAA
AAAGGAAGAGAGAGGAAGGGAGAAGGGAGAGGAATGAATTAGAGATGCCTCAGGCTTCAGA
CCTTACCTGGGTTTTCAGGGCAAACATAAATGTAAATACACTGATTTATTCTGTTACTAGATCAG
GTTTTAGGGTCCTGCAAAAGGCTAGCTCGGCACTACACTAGGGAATTTGCTCCTGTTCTGTCA
CTTGTCATGGTCTTTCTTGGTATTAAAGGCCACCATTTGCACAAATGTTCCTGTTTTGGGTAACTT
GGATTATTGTCAGGGGCCCgacgactgtagaaaagggccctgaagaagggccctctgctgtctagcGCTAGCG
CGGCCGCgaattc
